# A novel performance scoring quantification framework for stress test set-ups

**DOI:** 10.1101/2022.12.21.521346

**Authors:** Tal Kozlovski, Jeffrey M. Hausdorff, Ori Davidov, Nir Giladi, Anat Mirelman, Yoav Benjamini

## Abstract

Stress tests, e.g., the cardiac stress test, are standard clinical screening tools aimed to unmask clinical pathology. As such stress tests indirectly measure *physiological reserves*. The term reserve has been developed to account for the dis-junction, often observed, between pathology and clinical manifestation. It describes a physiological capacity that is utilized in demanding situations. However, developing a new and reliable stress test based screening tool is complex, prolonged, and relies extensively on domain knowledge. We propose a novel model-free machine–learning framework, the Stress Test Performance Scoring (STEPS) framework, to model expected performance in a stress test. A performance scoring function is trained with measures taken during the performance in a given task while exploiting information regarding the stress test set-up and subjects’ medical state. Multiple ways of aggregating performance scores at different stress levels are suggested and are examined with an extensive simulation study. When applied to a real-world data example, an AUC of 84.35[95%*CI* : 70.68 −95.13] was obtained for the STEPS framework to distinguish subjects with neurodegeneration from controls. In summary, STEPS improved screening by exploiting existing domain knowledge and state-of-the-art clinical measures. The STEPS framework can ease and speed up the production of new stress tests.

## Introduction

Stress tests, such as the cardiac stress test^1^, are commonly used in medical practice. They are especially useful when a physiological impairment is masked by a reserve mechanism. The term reserve describes a physiological capacity that may be utilized under stress, i.e., in demanding or emergency situations. This concept has been developed to account for the dis-junction, often observed, between pathology and clinical manifestation. The reserve concept is routinely used in several medical sub-fields, e.g., cardiac^1^, pulmonary^2^, and renal^3^.

These reserve related physiological mechanisms are often assessed by subjecting individuals to a “stress test” that compares function while the individual is at rest with that observed under exertion^4–6^. Thus, in stress tests a physiologic capacity is loaded by challenging a subject with tasks with an increasing levels of difficulty, consequently the functional reserve is increasingly loaded until it dysfunctions. Examples include pulmonary^2^, renal^3^, cardiac stress tests^1^. Stress tests have critical diagnostic and prognostic value. For example, reserve levels assessed via stress tests can dictate whether a transplant is a viable solution^7^ or a medical procedure is needed^8^. Stress test for other physiological systems such as the motor and cognitive reserve systems^9,10^ have yet to be introduced.

In the cardiac stress tests, the Bruce protocol^11^ is widely used for clinical assessment. Briefly, a treadmill is set to 1.7mph and at an inclined gradient of 10%. Every 3 minutes the incline of the treadmill is increased by 2%, and the speed increases as well. The test should be stopped when the subject cannot continue due to fatigue, pain or other medical condition. The test score is usually the maximal oxygen uptake (Vo2max). The Vo2max refers to the maximum amount of oxygen that an individual can take in and use during intense or maximal exercise. The Vo2max is approximated by a polynomial function of the test duration in minutes. Bruce et al.^12^ provided normal values of Vo2max that were estimated from a multistage treadmill testing procedure completed by healthy adults at various difficulty levels. Then, Bruce et al. presented nomograms for finding percent of deviation of an individual’s observed values of Vo2max from the normal values provided. During the years, the percent of deviations were refitted separately to different gender, BMI, age groups, and more.

Previously suggested stress tests, including the cardiac stress test, were developed by identifying a biological process and/or correlated measures to a latent reserve functionality. By understanding the reserve function mechanism and selecting a representative measure for this mechanism, clinical experiments revealed loadings levels that best predicted dysfunction^1,6^. Assessment of such prediction is done by comparing an individual’s function level with a reference function approximated with control subjects^13,14^. The distance, or percent of deviation, from the reference function can be viewed as a “performance score.” A Performance score that is below (or above) a set threshold indicates low reserve capacity.

The stress test modeling framework described above has certain disadvantages such as: (1) wide domain knowledge - researchers suggesting measures and loading levels for stress tests are based mainly on their knowledge and extensive trials, without benefiting from data driven methods. Data driven methods can help with revealing, or utilizing, high order interactions, which may be more difficult for a researcher to uncover. Moreover, such methods can help reduce the time required to unveil the best reserve predictors (the cardiac stress test took more than a decade to develop); (2) distributional assumptions - the use in classical statistical models, as regression and mean equality tests, often requires some assumptions on the data in-hand, such as Gaussian distribution or even completeness of the data collected. Usually, it is hard to predict what the data distribution will be and technical problems can occur that may interfere with data collection, and (3) lack of homogeneity in stress test modeling frameworks - different researchers are using slightly different methods to develop and/or describe the suggested stress test characteristics.

Generally, simple supervised classification models, such as logistic regression, are not suitable for predicting a reserve related impairment, mainly because the reserve mechanism helps to disguise such impairment. More specifically, a given dichotomy of healthy and impaired subjects will not be accurate due to the masking effect of the reserve mechanism. Thus, a longitudinal study is often required when developing a new clinical screening tool for a reserve deficiency. An extended follow-up helps to observe a significant deficiency in reserve levels that would change the subject medical status. Dyagilev et al.^15^ suggested the Disease Severity Score Learning (DSSL) framework that derives a disease severity score for a subject at a given time, from data based on clinical comparisons, i.e., pairs of disease states ordered by their severity. From those clinical comparisons, DSSL learns a function that maps the subject’s observed feature vector at time *t* to a severity score. Such a severity score would be a new scoring scale established using the DSSL framework. In order to establish such a new scoring scale, the objective function suggested in DSSL tends to maximize consistency of the subjects given medical severity ordering, while reducing high rates of temporal changes of the given scores. The DSSL is indeed useful, however, it only provides a one-dimensional line, as a function of time. For more complex cases, such as modeling stress tests, the objective function should account for another domain – the difficulty levels ordering.

Thus, based on the identified gaps we aimed to develop the Stress TEst Performance Scoring (STEPS) framework. The STEPS is designated for evaluating performance scores in stress tests. In such set-up, a subject faces some task with difficulty/loading levels that can be sorted by the difficulty level. The STEPS framework is aimed at learning a new function that would describe expected performance in a specific stress test design. To fulfill the required properties, we suggest extending the DSSL framework with an objective function that maximizes the distance of given scores for ranked pairs of subjects in a common difficulty level, and within-subject difficulty levels, while penalizing for high variance among repeated measures of the same task. We further suggest how to summarize performance scores to one indicative score to be used as a clinical screening tool for patients more likely to suffer from a reserve deficit.

## Methods

### STEPS Framework Overview

The goal of the proposed framework is to generate a scoring system that captures performance quality in challenging tasks. The performance score is a function of the observed feature vector collected during the stress test. Scores should further satisfy: (1) Between subjects concordance - performance scores should be correlated with the medical status, i.e., the performance of diagnosed individuals is expected to be poorer than that of healthy individuals (see figure 1a for illustration); (2) Within subject concordance - performance scores should be negatively correlated with a difficulty, i.e., performance degrades on challenging tasks) (see figure 1b for illustration); and (3) Low variance across repeated measures of the same individual at the same difficulty level (see figure 1c for illustration).

**Figure 1.**
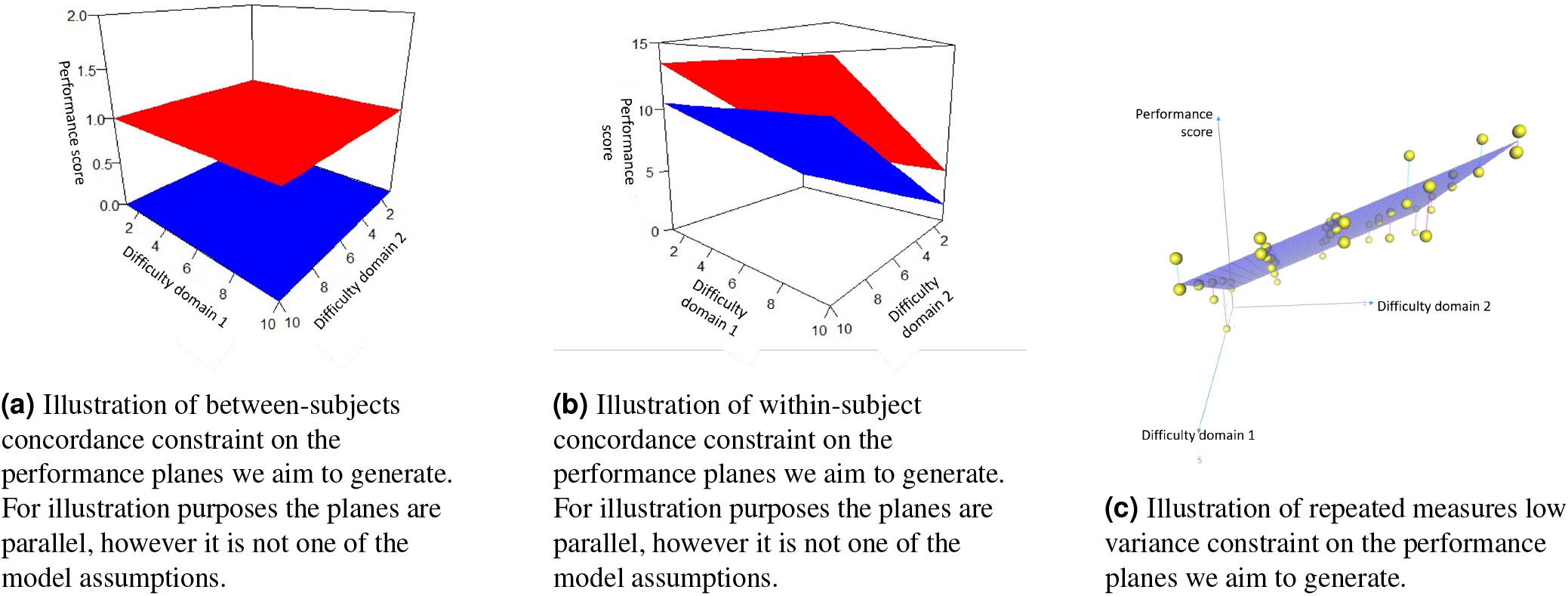
Illustrations of the constraints were imposed on the STEPS loss function.

Consider a feature vector, 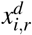, of subject *i, i* ∈ {1,…, *I*}, at difficulty level *d, d* ∈ *D*_*i*_, and repetition number *r*, 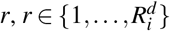, where 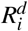 and *D*_*i*_ are the total number of repetitions subject *i* has of difficulty level *d* and the difficulties set of subject *i*, respectively. This feature vector holds information collected during a task completeness (e.g., heart rate during a cardiac stress test). This information should be summarized into a single performance score, representing the performance in the challenge level *d*. This score is produced by learning a function 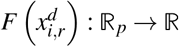 for a given feature vector, 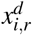. The score will impose clinical ordering of subjects according to their clinical state using a set of ordered pairs of subjects, *B*, as well as imposing challenges difficulty ordering using a set of ordered pairs of difficulty levels subject *i* encountered, *W*_*i*_. Furthermore, a set of repetitions of subject *i* at level *d*, 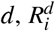, would help to reduce variability among repeated measures of the same task. A representative performance score of subject *i* at difficulty level *d* would be the average across given scores of all her repetitions, 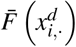. To produce performance scores that satisfies the required properties as detailed above, an optimization problem is solved to minimize an additive loss function. The loss function is composed of three terms: the between-subjects term, the within-subjects term, and the repeated measures term.

From the STEPS produced scores, one can portray subjects’ performance plane, spanned by possibly more than one dimension, as a function of the stress level encountered. The resulted functions of the control subjects in the training set will be used to construct a reference function. An aggregated score is produced according to the distance of an individual’s performance plane to the reference function. The distance value per subject can be used in a classification model, upon which an appropriate threshold for indicating low reserve capacity will be determined.

#### Between Subjects Term

The first requirement from the STEPS function is to distinguish between clinical states of subjects, i.e., the produced score should maximize the consistency of the severity ordering in each challenge (or in a specific subset of challenges according to the stress test design). The between subjects term (Equation 1), *L*_*B*_, utilizes existing domain knowledge.

Let *B* be the set of between subjects ordered pairs according to their disease severity. Set *B* includes pairs of ordered subjects ⟨*i, j*⟩ indicating that subject *i* is in a more severe state than subject *j*. |*B*| is the number of ranked pairs (similarly, we will use |·| to specify the size of a set). Set *B* is generated according to clinical knowledge. Next, each pair ⟨*i, j*⟩ is associated with a set *D*_ij_, that holds all common challenges both subjects encountered (or a subset of challenges that we want to consider through the STEPS framework learning stage).

The objective function is constrained to maximize consistency of the severity ordering, that means that the learned STEPS function should produce a higher score for subject *i* at difficulty level *d* than for subject *j*, by penalizing when this requirement does not hold. This penalization for an ordered pair *i j* in difficulty level *d* is induced by using the Huber approximation to the Hinge loss function^16^: 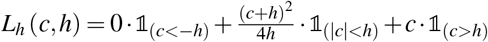. Where, for our purpose 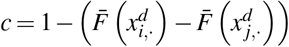, thus for small differences in scoring and/or for opposite ordering, there is a growing penalty in the scores difference. The between subjects term is the average penalty of an ordered tuple ⟨*i, j*⟩, across all their common difficulties *D*_*i, j*_, and a further average is done across all tuples in set *B*:

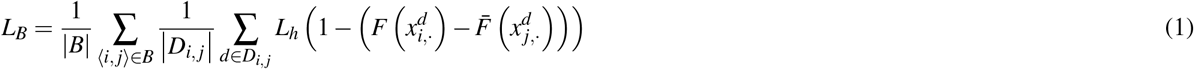

#### Within Subjects Term

The second requirement is to distinguish between difficulty levels without using the information of the difficulty level as part of 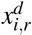. A stress test is designed to increase loading levels until a physiological reserve fails, also referred as the “tipping point” in which performance starts to decline as compared to normal values. This point is different between subjects, and conceptually it reflects the anticipated reserve decline. Therefore, the STEPS framework should yield a score that increases within subject’ records according to the difficulty encountered, i.e., better performance is expected in easier challenges. Therefore, removing the difficulty information from the feature vector at hand would loosen our constraint and give the learning process the freedom to find differences in performance according to collected variables.

Let *W* be the set of within-subjects ordered pairs of difficulties of the form ⟨*d, d*^′^⟩, where *d*^′^ is the easier difficulty level. Performance at level *d*^′^ is expected to receive a lower score than in *d*, reflecting the better performance in the task. *W*_*i*_ is the subset belonging to subject *i*, and |*W*_*i*_| is the number of ordered pairs belongs to subject *i*. Similar to the between subjects term, the within subjects term (Equation 2), *L*_*W*_, penalizes differences between scores of the same subject in different difficulties. Furthermore, the within-subjects term is the average penalization within a subject, and a further average of penalties across all subjects, as follows:

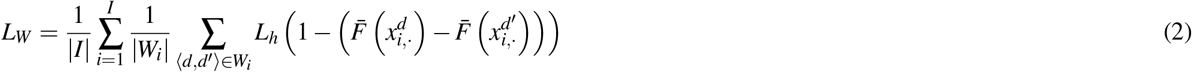

#### Repeated Measures Term

The last requirement from the STEPS function is to be consistent with within-subject challenge repetitions. For that reason, a variance term of the repeated measures of a subject was added to the objective. The expectation is that each subject will score similarly across identical difficulty levels.

Let 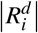 be a set that holds the repeated measures of subject *i* in a difficulty level *d*. Furthermore, |*D*_*i*_| is the number of unique difficulties subject *i* faced, and 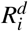 is the number of repetitions of subject *i* in difficulty *d*. In order to penalize large discrepancies among repeated measures, this term (Equation 3), *L*_*R*_, is defined to be the average variance of the fitted performance scores in each set 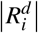, and a further average is done across all subjects:

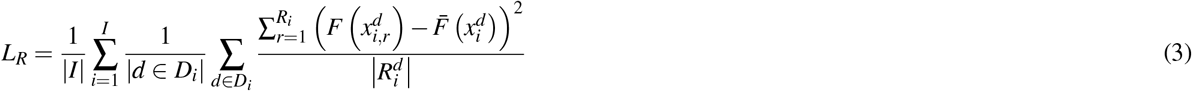

#### Objective Function and Optimization

By summing the three terms, and adding coefficients, *λ*_*W*_ and *λ*_*R*_, that control the within-subjects and the repeated measures terms’ relative weights to the between-subjects term (coefficients values can be determined through cross-validation), we receive the following convex and twice differentiable objective function (see Appendix A for derivatives): 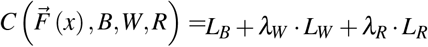

To train the STEPS function, we seek to minimize the objective function using Gradient Boosting Regression Trees (GBRT) algorithm^17^, where shallow greedy regression trees (implemented using the rpart package^18^) are used as weak learners^19^. See algorithm 1 for more details on the learning stage of the STEPS function *F*.

##### Algorithm 1

GBRT algorithm to learn a STEPS function

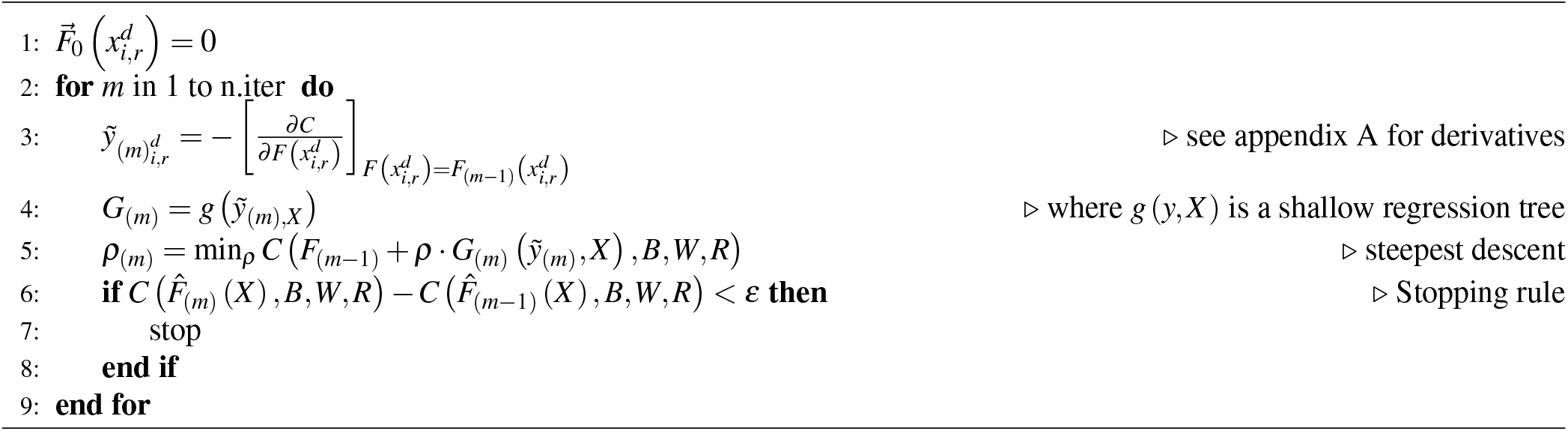

#### Other Optimization Methods and Code Availability

The STEPS as presented in this paper focuses on evaluating a non-linear function. As was detailed in Dyagilev et al.^15^, other optimization methods can be used. In addition, one can assume *F* (*x*) is a weighted linear combination of the feature vector, i.e. 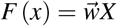, and optimize the objective function with a Newton-Raphson numeric search. Both linear and non-linear optimization methods were implemented in the STEPS R package, and are accessible at: https://github.com/TalKozlovski/STEPS/tree/main/STEPS

### Aggregation to performance score index

Algorithm 1 scores the performance in each challenge a subject encountered during a stress test. In order to quantify reserve capacity, a subject’s performance scores need to be aggregated across all difficulties encountered into one indicative index that reflects the subject’s reserve capacity. This index can be obtained by quantifying the distance of a subject’s performance plane, i.e. performance score as a function of difficulty level, from a reference performance plane. The reference performance plane is the median performance score among the healthy subjects used for training the GBRT model. This reference function is expected to be lower than the similarly constructed plane of subjects diagnosed with reserve dysfunction. Thus, if a performance plane of one subject is closer to the reference function than that of another subject, it would indicate the former has a higher reserve capacity than the latter. There are multiple ways to measure the distance from the reference plane, as detailed below. Their performance assessment is reviewed through simulation at the next section. The suggested aggregation methods that we examined:

1. Mean difference from the reference plane across all difficulty levels.
2. Trimmed mean difference from the reference plane - same as above except for outliers exclusion.
3. Mean of positive differences from the reference plane - since higher scores indicate worse performance, there is a logic in averaging distances on scores that were higher than the median of the control group.
4. Mean of SD-positive differences - average positive differences that are farther than 1 unit of SD from the reference plane.

### Simulation study

To demonstrate the STEPS framework (Figure 2) feasibility, a simulation study was conducted. For that purpose, consider a synthetic clinical state that is known to be masked with an associated physical reserve. Therefore, under stress, it is expected that performance will decline or reach the “tipping point”. A subject is considered positive if her sampled severity is higher than the median of all subjects. Thus, our interest lies in discovering subjects exceeding this threshold. Further assume that severity level is affected by two independent domains, *s*_1_ and *s*_2_, each sampled from *N* (10, 1) (mean is 10 to avoid negative values). The severity is computed as their multiplication, *S* = *s*_1_ · *s*_2_. Moreover, there are 2 available clinical measures used to estimate *S, score*1 and *score*2, each dependent in one of the severity domains. Namely, *score*1 = *s*_1_ + *ε* and *score*2 = *s*_2_ + *ε*, where *ε*∼ *N* (0, 1). *score*1 and *score*2 represents the existent domain knowledge, which will be utilized with the STEPS framework to score the simulated stress test.

**Figure 2.**
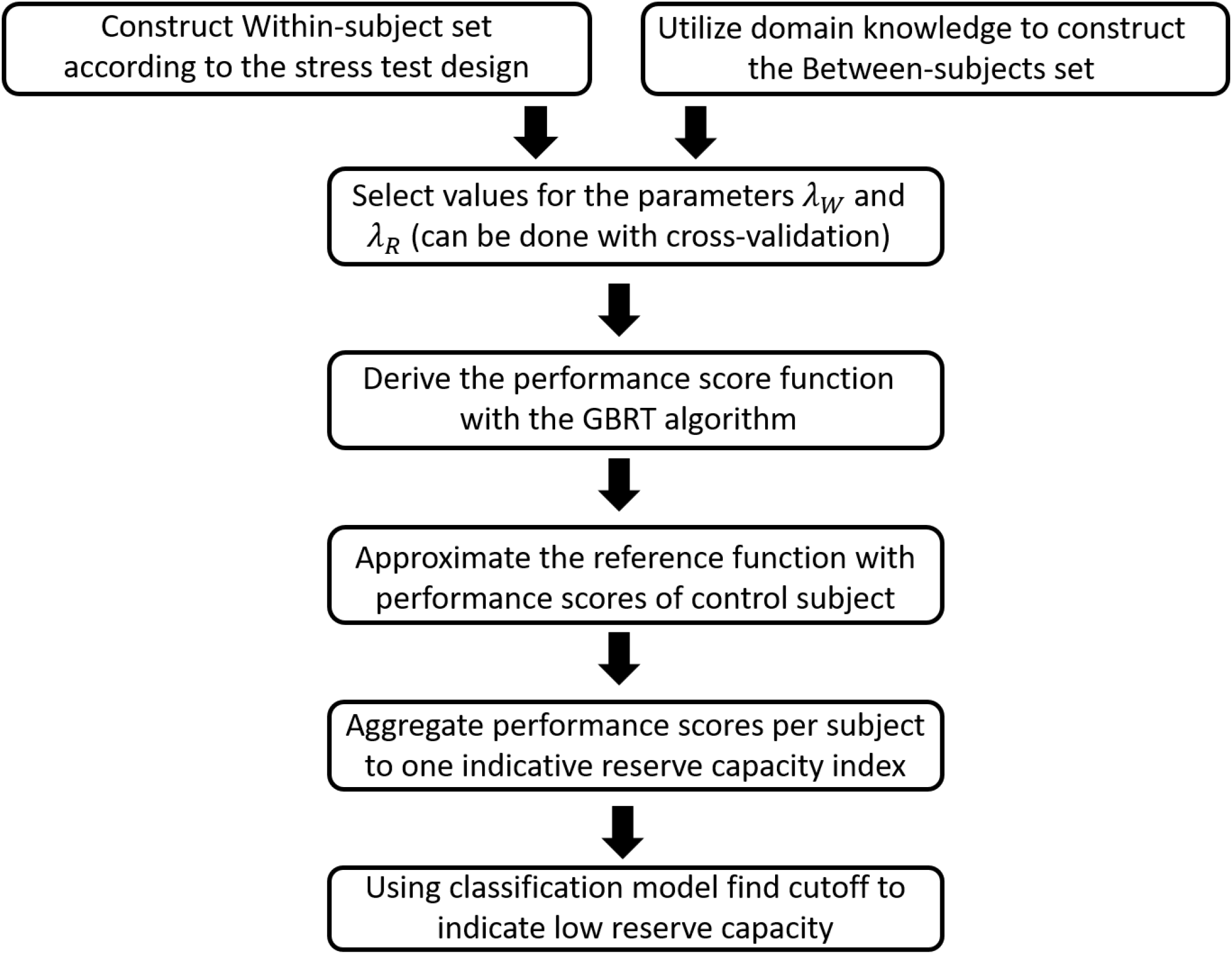
STEPS framework

The simulated stress test consists of 2 difficulty domains *d* ∈ {*d*1 × *d*2}, each includes 3 difficulty levels, *d*1, *d*2 ∈ {0, 1, 2}, and 2 repetitions for each subject in each difficulty level, 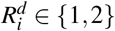. A feature vector of subject *i* at repetition *r* of difficulty level 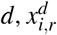, holds 5 entries. The signal is concentrated in one entry and is a function of both *S* and difficulty level (we will refer to this measure as the observed severity – Figure 3). The other four entries contains only noise. The noise level is simulated from *N* (0, *σ*) ^2^, where *σ* ^2^ varies between 1 to 0.01. This range is used because we assume that constructing a stress test should improve the use of available measures (*score*1 and *score*2), which here are simulated with *ε* ∼ *N* (0, 1).

**Figure 3.**
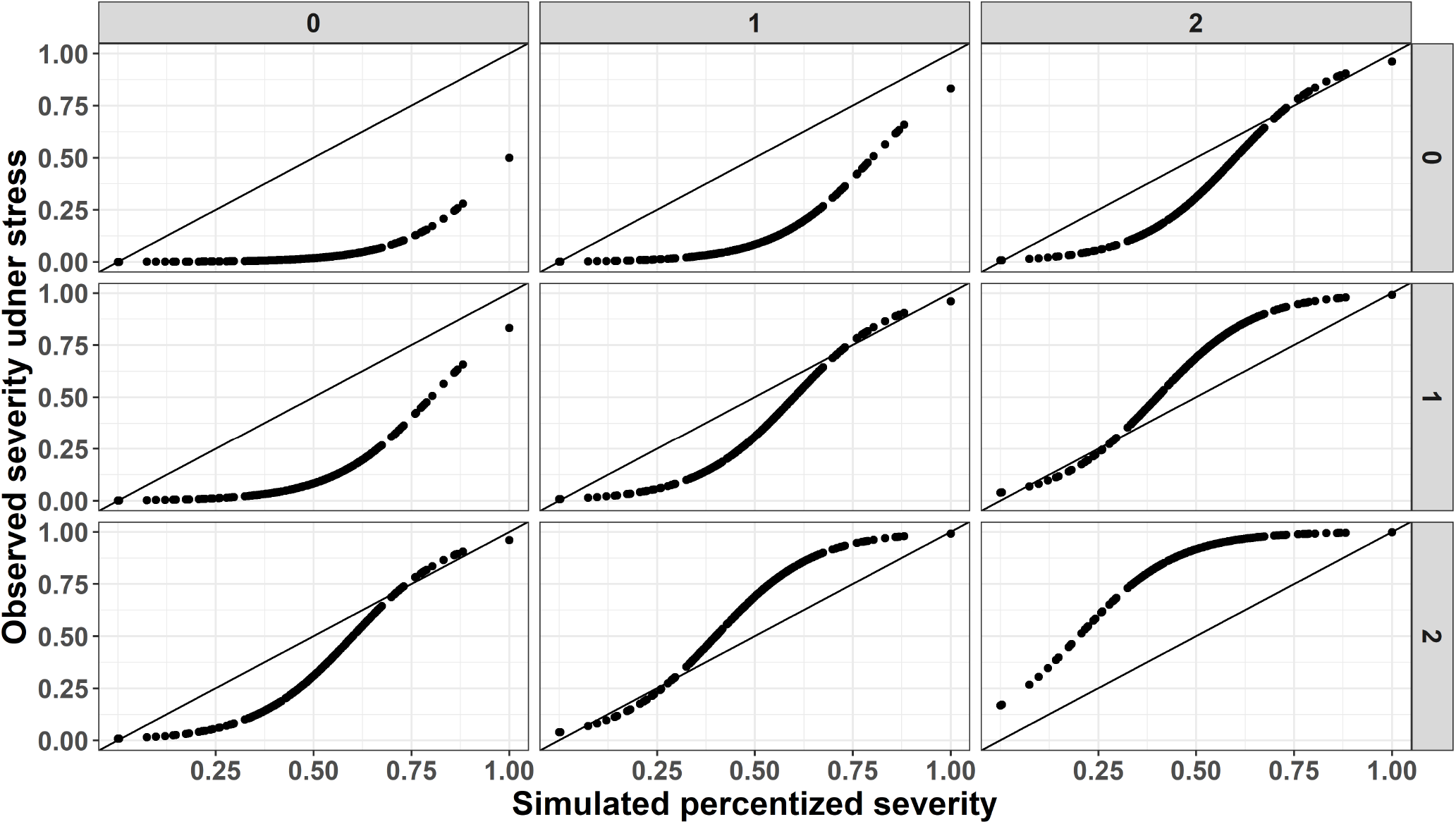
Illustration of simulated severity versus the observed severity as simulated from conducting a stress test. Columns facets presents difficulty levels of domain # 1 and rows facets presents the levels of difficulty domain # 2.The black solid lines are identity lines.

The idea behind simulating the observed severity is that a stress test should extend the physiologic reserve in question to its limit, and therefore help reveal the true underlying severity state of a subject. According to the above, the observed severity is a function of *S* and difficulty levels the subject had encountered, where we assume that in easy difficulty levels the observed severity is good for most subjects (excluding the most severe cases). As the challenges increases, even the low severity cases will yield deteriorated performance, similarly to those with higher severity to begin with. To do as noted, *S* was set to receive values between (0, 1), and a logistic transformation, 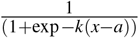 was applied, with a fixed growth rate (*k*) of 8, and decreasing sigmoid’s midpoint (a) as a function of the sum of the two difficulty domains (i.e. midpoint is 1 for sum difficulties of 0, 0.8 for sum of 1 and so on). Figure 3 presents the simulated observed severity as a function of severity and difficulty levels.

The sets required for the STEPS framework were constructed as follows: for the Oracle model, set *B* was based naively on *S*, and otherwise according to *score*1 and *score*2 (if subject *i* has both values higher than 1 SD unit from subject *j* than a pair is considered ordered). Set *W* was constructed by holding fixed one difficulty dimension and comparing the other difficulty dimension levels, and vice versa. The within subjects set has 18 comparisons per subject. The STEPS function using only the 5 stress test features as covariates.

The simulation includes 200 subjects, with an 80% - 20% train-test sets division. Further required STEPS parameters that were used are 50 trees (or convergence rate of 10^−4^), and GBRT step size is 1 and not optimized to enhance calculations. Last but not least, we set *λ*_*B*_ and *λ*_*R*_ to 1 and varied *λ*_*W*_ respectively between values of 0.01; 0.5; and 1. Each scenario was repeated 500 times. The simulation script, and a summary report can be found at: https://github.com/TalKozlovski/STEPS/tree/main/STEPS_simulation

### Simulation results

The simulation output is reviewed with the following summary measures: (1) Concordance values of STEPS and Oracle-STEPS in reconstructing the ordered sets *B* and *W*, i.e., the percent of correctly reconstructed pairs according to their given predictions, and average SD values for set *R*; (2) Correlations between severity, *S*, to aggregated scores of STEPS, and Oracle-STEPS (aggregated scores as presented earlier), as well as to *score*1 and *score*2; and (3) Area Under the Curve (AUC) values to estimate classification abilities of aggregated scores of STEPS, Oracle-STEPS, as well as *score*1 and *score*2. The STEPS framework uses learning sets of pairs computed according to values observed in *score*1 and *score*2. Thus, we expect that, at a similar noise level, the STEPS output would be at most as good as using *score*1 and *score*2, and will outperform *score*1 and *score*2 as noise levels decreases.

#### Concordance values

First, we examine the concordance levels achieved in reconstructing sets *B* and *W* (Figure 4). Overall, the observed STEPS model showed higher SD values than the Oracle model. Concordance values becomes better with the increase in noise ratio, and for noise ratio of 1 concordance levels are around 0.5 for both sets. As *λ*_*W*_ increases the STEPS gets closer results to the Oracle-STEPS model. Concordance of set *B* converges to 0.8. Concordance of set *W* converges to 0.9. Second, the average SD values among set *R* are surprisingly higher for the Oracle-STEPS. This can be explained due to relatively better achievements in reordering sets *B* and *W* which probably masked the necessity to improve prediction within the repeated measures term that takes in account set *R*. Both average SD values are much smaller than the loss function parameter used, *h* = 0.5.

**Figure 4.**
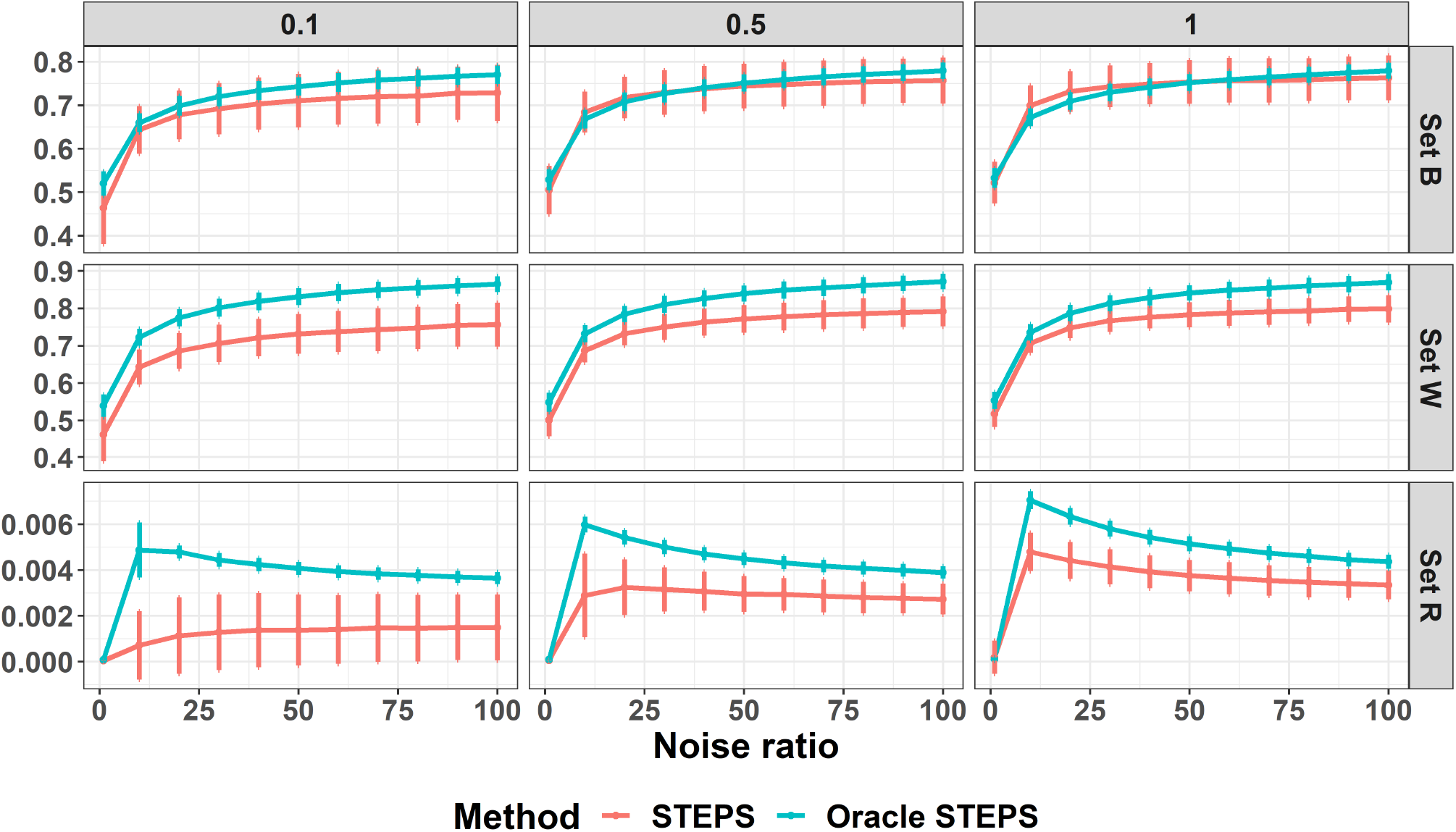
Concordance levels for reconstruction of sets *B* and *W*, and average SD values for set *R*. X-axis presents noise ratio of that added to stress test features relative to the noise added to the observed clinical scores. Y-axis presents concordance for sets *B & W* and SD values for set *R*. Rows are for different sets and columns presents the different *λ*_*W*_ values used in simulation.

#### Correlation to latent severity

Since the goal in using the STEPS framework is to estimate a latent severity that reflects reserve capacity and may be masked, it is important to review the correlation of given scores to the severity measure, *S*. Aggregated scores of trimmed mean difference from median returned equal scores to that of mean difference from median (probably due to the normal like distribution of the produced scores). On the other hand, Mean of SD positive difference from median had very poor achievements and therefore are no longer reviewed. In Figure 5, the correlations between produced scores and *S* are presented. The latent severity is linearly dependent on *score*1 and *score*2. Thus, as expected the correlation between *S* and both scores is ∼0.5. When noise ratio is 1 all models based on the STEPS framework have the lowest correlation with *S*, and as the noise ratio increases the STEPS framework reached Spearman’ correlation levels of ∼0.8. The mean difference from median performs better than the mean positive difference from median. Again, the increase in *λ*_*W*_ helps to narrow the gap in correlation levels between STEPS and Oracle-STEPS.

**Figure 5.**
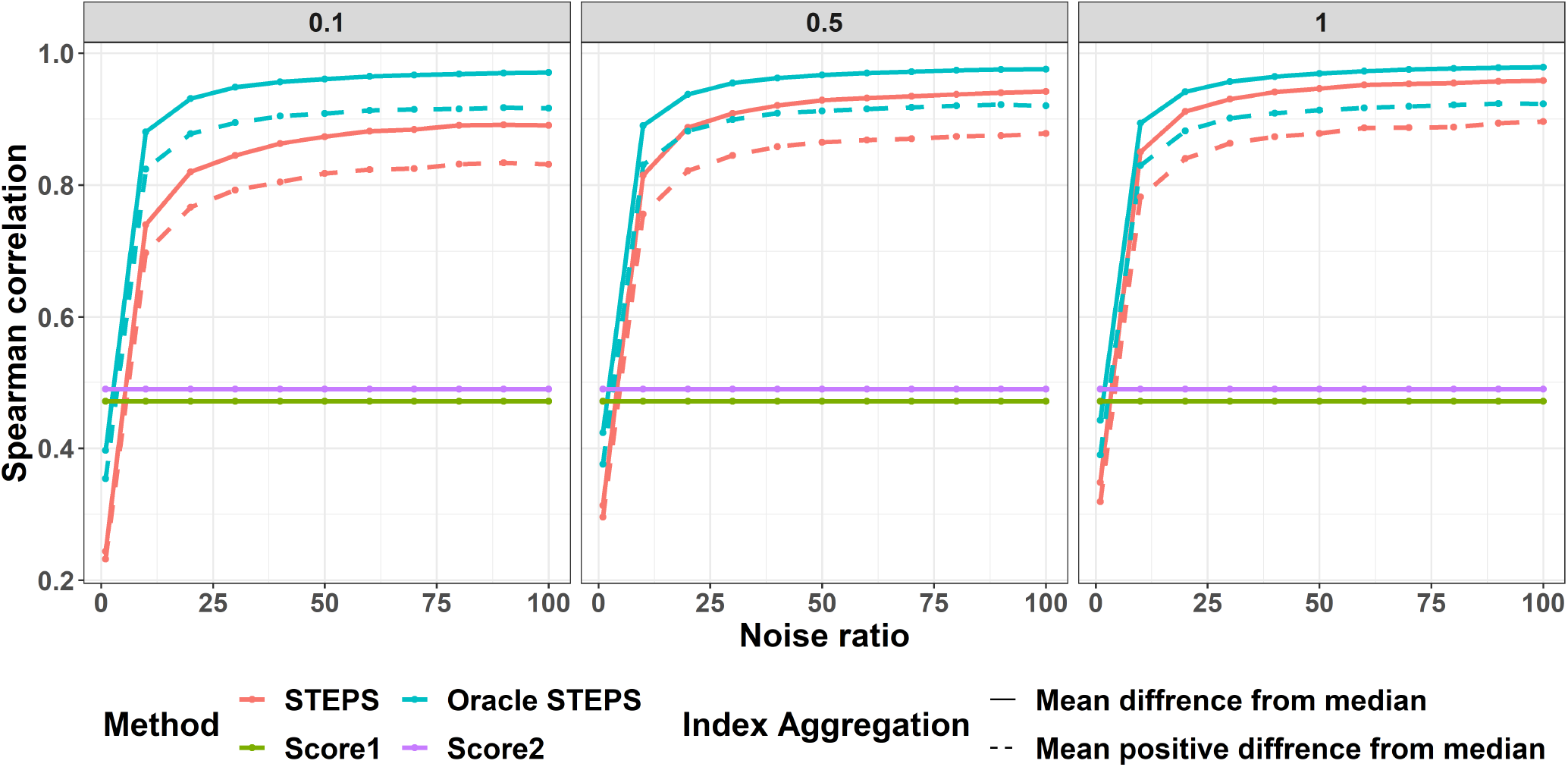
Correlations between all scores to the latent severity. X-axis is the noise ratio. Y-axis Spearman’s correlation coefficient. Color represents the different models and line type distinguishes between aggregation methods for the STEPS scores. Columns are different *λ*_*W*_ rates.

#### Classification model

In regards to AUC values (Figure 6, both aggregated scores, mean difference from median and mean of positive differences from median, showed similar behaviors. As expected, at noise ratio of 1, the Oracle-STEPS reaches similar AUC values to that of *score*1 and *score*2 (∼0.72). As the noise ratio increases, the STEPS based models outperforms the use of *score*1 and *score*2 for classification. Moreover, the use of higher *λ*_*W*_ helps to reach AUC levels closer to 1 and the convergence is faster (in terms of noise ratio).

**Figure 6.**
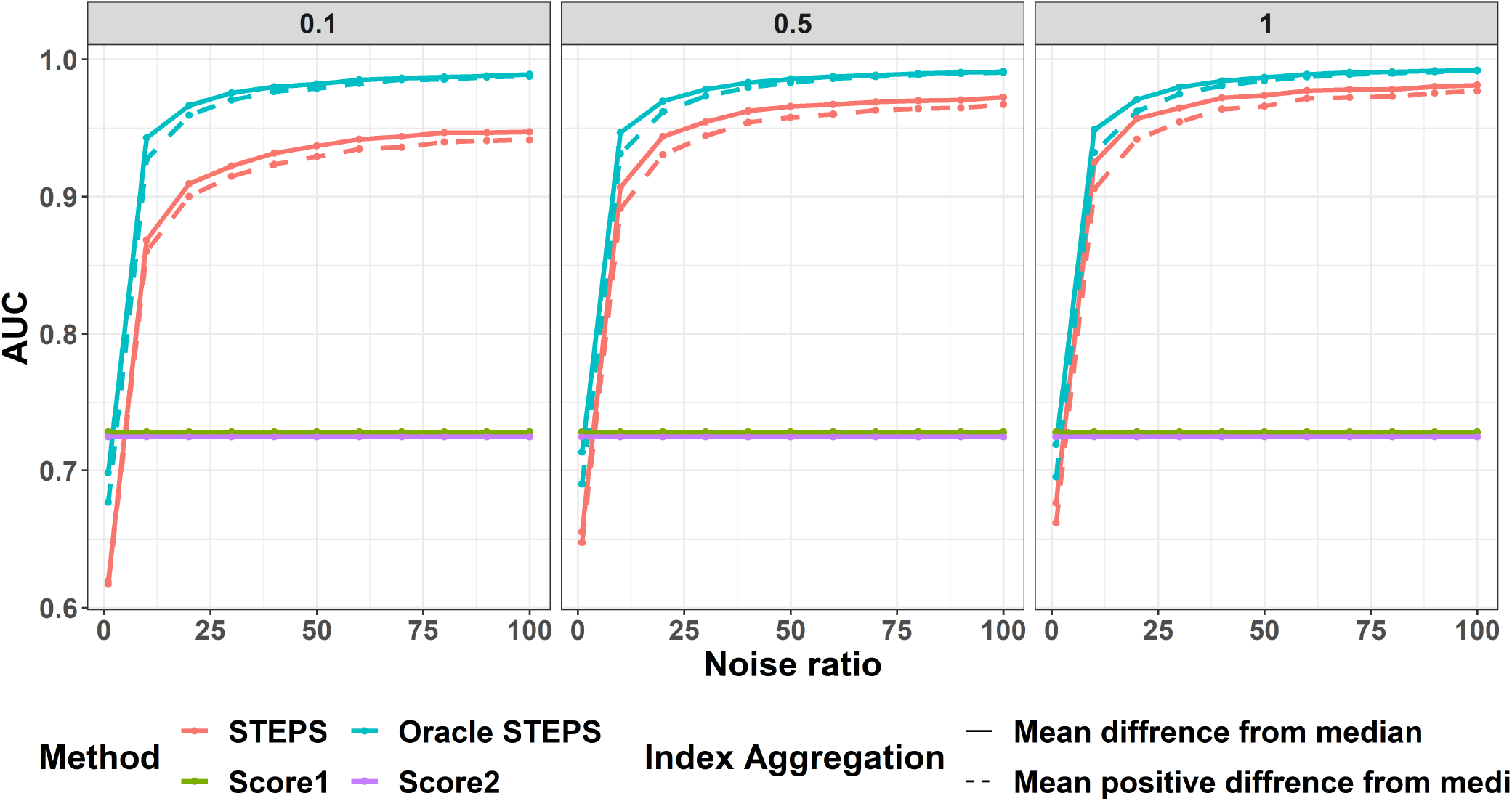
AUC values. X-axis is the noise ratio. Y-axis average AUC values. Color represents the different models and line type distinguishes between aggregation methods for the STEPS scores. Columns are different *lambda*_*W*_ rates.

### Real data example

In this section, the STEPS framework and two of the suggested score aggregation methods (mean difference and trimmed mean, as those performed best in the above simulation study) are illustrated using a clinical dataset. To explore the utility of the STEPS framework we wanted to examine a multi-layer, multi-dimensional task that could account for more than one dimension.

In the past decade, there has been a shift in the understanding that functional abilities are likely to be multi-factorial processes that rely on both motor and cognitive function. Given the growing global burden of dementia, there is an urgent need to improve the clinical identification of older persons at risk for dementia. The dependency between cognitive and gait dysfunction has been found even in otherwise healthy older adults with subtle cognitive deficits that are not yet profound enough to be identified using global measures of cognitive status^20^. From a practical perspective, this provocative time course suggests that it may be possible to augment the prediction of cognitive decline and mobility based on objective gait evaluation, under challenging conditions, if so, this presents an early opportunity for appropriate interventions.

Data were collected at the Laboratory for Early Markers of Neurodegeneration (LEMON) at Tel Aviv Medical Center. The study was approved by the Tel Aviv Medical Center Ethics Committee (635-16-TLV and NCT01730599). All subjects provided informed written consent for the study according to the Declaration of Helsinki 1964. 156 patients diagnosed with Parkinson’s disease (PD) and 274 healthy controls were recruited (Table 1). The dataset includes features collected during a clinical gait assessment, composed of usual-walking and dual task (serial subtraction) walking, a paradigm used to assess gait dysfunction in PD^21^.

**Table 1.** Mean (SD) of patients characteristics, and the number of males (%), per diagnosis group.

|  | Control | PD |
| --- | --- | --- |
| Age | 52.7 (10.66) | 64.24 (9.17) |
| Male n ( % ) | 113 (41.24%) | 99 (63.46%) |
| Disease duration (years) | NA | 2.74 (2.32) |
| Years of education | 17.45 (2.62) | 16.56 (2.93) |
| Preferred usual walking speed (m/s) | 1.37 (0.22) | 1.18 (0.26) |
| Preferred dual-task walking speed (m/s) | 1.18 (0.26) | 0.98 (0.25) |
| MOCA | 26.45 (2.7) | 23.86 (3.81) |
| TMTA (sec) | 41.71 (16.41) | 71.51 (42.02) |
| TMTB (sec) | 78.54 (29.04) | 130.31 (70.95) |

PD is a neurodegenerative disease with a long prodromal phase during which signs and symptoms are not visible^22^. As such, PD patients in the prodromal and early stages of the disease may compensate for the accumulated damage and perform at a level comparable to a normal population. The compensation mechanism is often enabled by what is referred to as brain or cognitive reserve^23^. This reserve functionality can be stressed by enhancing brain activity demands, for example, by challenging a patient with a dual-task, composed from motor execution in parallel to a cognitively challenging task^24^.

The clinical gait assessment included walking along a 20-m-long corridor under two conditions: (1) preferred, usual-walking speed, and (2) dual-tasking, serial 7 subtraction task starting from a three-digits number^25^. Another aspect of this assessment is the Timed Up and Go test (TUG)^26^. The TUG test consisted of standing up from a chair, walking 3 m to a designated location at a normal pace, turning around, walking back, and sitting back down on the same chair. Using small three-axis accelerometers with a gyroscopes worn on the lower back and sensors located on both ankles, in-lab gait features (e.g., gait speed, step regularity, and symmetry measures) were quantified as previously described^27^,^28^.

In parallel to the clinical gait assessment, the participants completed two cognitive tests: the Montreal Cognitive Assessment – MOCA^29^ which is a measure of general cognition and the Trail Making Test-TMT^30^ which is a test of scanning (Section A) and executive function (Section B). Participants were also assessed with their preferred walking speed during a usual walk and during the dual-task walk to evaluate motor function. Those tests represent the domain knowledge and are of assistance in creating the between-subjects ordered set. The within-subjects set is constructed by pairs of instances of dual-task vs. usual walk of the same subject. Each subject completed each task once. Thus, the repetitions set weight parameter, *λ*_*R*_, is set to 0. Moreover, following our simulation results, *λ*_*W*_ and *λ*_*B*_ were set to 1.

The dataset was divided into 10-folds, where the train set is used to construct the STEPS model and to fitting logistic regressions on the STEPS aggregated scores or clinical features. The test set kept aside is used only to estimate AUC levels, and to construct 95% bootstrap CI’s. AUC values and CI’s were obtained from the ‘pROC’ R package^32^ with 2000 bootstrap samples within each fold. The AUC curve obtained in each fold for each of the measures, was further tested against the Mean-STEPS difference measure’s AUC curve for a significant difference. Each measure yielded 10 such pvalues, where the respective minimum and a combined Harmonic Mean P-value (HMP)^33^ are provided for.

Results show (Table 2) that the two STEPS scores’ aggregation methods have better AUC levels than all the clinical measures used for clinical screening. In this task, we aimed to use the STEPS aggregated scores to be used as a classification tool. The classification tool should identify diagnosed patients and help identify the non-diagnosed that have a higher probability of manifesting in the future. Future studies should explore performance in prodromal screening with the utility of STEPS in cohorts of individuals at risk for developing neurodegenerative conditions like Alzheimer’s disease and Parkinson’s disease.

**Table 2.** Average AUC levels and average bounds of 95% bootstrap CI’s across 10-folds for distinguishing patients according to given medical status. AUC levels were tested between each measure and the STEPS-Mean difference measure within each fold, with the DeLong test. Minimal p-value, as well as the Harmonic Mean P-value (HMP) are presented.

| Measure | AUC; [95% CI] | Min P-value | HMP |
| --- | --- | --- | --- |
| STEPS-Mean difference | 84.35; [70.68, 95.13] | NA | NA |
| STEPS-Trimmed mean difference | 84.35; [70.41, 95.13] | 1 | 1 |
| TMTA (sec) | 78.65; [62.49, 91.61] | 0.022 | 0.120 |
| TMTB (sec) | 78.05; [62.07, 91.24] | 0.004 | 0.036 |
| Average dual-task walking speed | 73.43[56.03, 88.35] | 0.009 | 0.042 |
| Average usual walking speed | 71.84[54.90, 86.56] | 0.001 | 0.010 |
| MOCA | 69.91; [52.03, 85.61] | < 0.001 | 0.002 |

## Discussion

This paper presents the novel Stress TEst Performance Scoring (STEPS) framework for stress test set-ups. The STEPS automates the development of stress tests by incorporating domain knowledge and machine learning. While the reserve is unobservable, a domain expert can still rank (partially) subjects according to their medical records or other clinical assessments. The ranking is used to fit a semi-supervised model using the data collected during the stress test. The model assigns each subject a score that best agrees with the domain expert ranking. The score serves as the proxy of the reserve. Using a linear function as our model will yield an interpretable explanation of the reserve. While using GBM captures the non-linear patterns in data, resulting in a more accurate model. The STEP framework is applicable when using partial data (partial in terms of difficulty measures, i.e., subjects do not require to complete the whole stress test if allocated to the train set).

The STEPS framework was tested using a simulation study and demonstrated on real data. The simulation showed the advantage of using the framework to exploit existing domain knowledge in a machine learning model to generate a new scoring system. Thus, given that a suggested stress test can better measure a latent reserve capacity than existing clinical classification tools, it can be more beneficial in estimating real latent severity. I.e., reserve capacity decline and be more accurate when further used for classification.

Specifically, we demonstrate the STEPS framework on a task to quantify motor and cognitive reserve deterioration among PD subjects. We interpreted the already clinically used Timed Up and Go test as a challenging stress test. The difficulty levels in this assessment are an easy challenge of a typical walk and a more challenging level of a dual-task walk. During these challenges, gait features were collected. The other clinical tests were used to rank the subjects according to their severity. Applying the STEPS framework results in a new scoring system that outperformed common clinical measures from both the cognitive and motor domains.

A limitation of the real data example is that we are measuring the utility of the various methods in the accuracy of classification of the subjects to healthy and afflicted. However, our goal is to quantify motor-cognitive reserve. Unfortunately, the reserve quantification can only be validated in a longitudinal study, where the subjects are tracked for a manifestation of a neurodegeneration.

There are several avenues for future work. Besides applying the current STEP framework in a longitudinal study and validating the reserve quantification, several theoretical advances exist to consider. The procedure can be modified to adapt to the subject’s performance in the test. Adjusting the difficulty level not to frustrate or bore the patient while ensuring sufficient data is collected. Additionally, developing methods to estimate the relation between the various features collected in the stress test and the score performance. An interesting approach is to use features importance as in^17^, SHAP values^34^ or statistical testing.

In summary, this paper presents a novel modeling tool for physiological stress test set-ups. Stress tests are common diagnosis tools for reserve deficits. The model outputs a score that reflects a subject’s performance in the stress test. Under the assumption the stress test is rightly targeting the subejct’s reserve functionality, then this score can imply the subject’s reserve capacity. The model leverages on clinical knowledge to model performance in a stress test set-up. The rationale behind stress tests is to model the compensation mechanism, not the observed medical state. Furthermore, although only given as an example, it is the first time a stress test is suggested to quantify the brain reserve mechanism, which is known to have a complicated compensation mechanism.

## Acknowledgements

The authors would like to thank all the participants in the clinical gait assessment and the LEMON staff for their assistance with the data collection. The authors also would like to thank Tzviel Frostig for his help regarding the computational challenges in the simulation study, and his input.

## Author contributions statement

T.K. conceived and developed the STEPS algorithm, conducted the experiments and the real data analysis, and wrote the main draft of the manuscript, A.M., and Y.B. guided direction of research, interpreted the results, and revised the manuscript, O.D.,

N.G. and J.H. interpreted the results, and edited the manuscript.

## Additional information

**Accession codes** https://github.com/TalKozlovski/STEPS/tree/main/STEPS

## References

1. Cooke, G. et al. Physiological cardiac reserve: development of a non-invasive method and first estimates in man. Heart 79, 289–294 (1998).

2. Gorcsan III, J., Murali, S., Counihan, P. J., Mandarino, W. A. & Kormos, R. L. Right ventricular performance and contractile reserve in patients with severe heart failure: assessment by pressure-area relations and association with outcome. Circulation 94, 3190–3197 (1996).

3. Bosch, J. P. et al. Renal functional reserve in humans: effect of protein intake on glomerular filtration rate. The Am. journal medicine 75, 943–950 (1983).

4. Biagini, A. et al. Early assessment of coronary reserve after bypass surgery by dipyridamole transesophageal echocardiographic stress test. Am. heart journal 120, 1097–1101 (1990).

5. Gibby, C., Wiktor, D. M., Burgess, M., Kusunose, K. & Marwick, T. H. Quantitation of the diastolic stress test: filling pressure vs. diastolic reserve. Eur. Hear. Journal–Cardiovascular Imaging 14, 223–227 (2013).

6. Sharma, A. et al. Optimizing a kidney stress test to evaluate renal functional reserve. Clin. nephrology 86, 18 (2016).

7. Ader, J.-L. et al. Renal functional reserve in cyclosporin-treated recipients of kidney transplant. Kidney international 45, 1657–1667 (1994).

8. Blomqvist, C. G. Use of exercise testing for diagnostic and functional evaluation of patients with arteriosclerotic heart disease. Circulation 44, 1120–1136 (1971).

9. Stern, Y. Cognitive reserve. Neuropsychologia 47, 2015 (2009).

10. Elbaz, A. et al. Motor function in the elderly: evidence for the reserve hypothesis. Neurology 81, 417–426 (2013).

11. Bruce, R., Blackmon, J., Jones, J. & Strait, G. Exercising testing in adult normal subjects and cardiac patients. Pediatrics 32, 742–756 (1963).

12. Bruce, R., Kusumi, F. & Hosmer, D. Maximal oxygen intake and nomographic assessment of functional aerobic impairment in cardiovascular disease. Am. heart journal 85, 546–562 (1973).

13. Tan, L. Cardiac pumping capability and prognosis in heart failure. The Lancet 328, 1360–1363 (1986).

14. Tan, L.-B. & Littler, W. A. Measurement of cardiac reserve in cardiogenic shock: implications for prognosis and management. Heart 64, 121–128 (1990).

15. Dyagilev, K. & Saria, S. Learning (predictive) risk scores in the presence of censoring due to interventions. Mach. Learn. 102, 323–348 (2016).

16. Rennie, J. D. & Srebro, N. Loss functions for preference levels: Regression with discrete ordered labels. In Proceedings of the IJCAI multidisciplinary workshop on advances in preference handling, vol. 1 (Kluwer Norwell, MA, 2005).

17. Friedman, J. H. Greedy function approximation: a gradient boosting machine. Annals statistics 1189–1232 (2001).

18. Therneau, T., Atkinson, B., Ripley, B. & Ripley, M. B. Package ‘rpart’. Available online: cran. ma. ic. ac. uk/web/packages/rpart/rpart. pdf (accessed on 20 April 2016) (2015).

19. Zheng, Z. et al. A general boosting method and its application to learning ranking functions for web search. Adv. Neural Inf. Process. Syst. 20: Proc. 2007 Conf. (2007).

20. Montine, T. J. et al. Concepts for brain aging: resistance, resilience, reserve, and compensation. Alzheimer’s research & therapy 11, 1–3 (2019).

21. Mirelman, A. et al. Gait impairments in parkinson’s disease. The Lancet Neurol. 18, 697–708 (2019).

22. Schrag, A., Anastasiou, Z., Ambler, G., Noyce, A. & Walters, K. Predicting diagnosis of parkinson’s disease: a risk algorithm based on primary care presentations. Mov. Disord. 34, 480–486 (2019).

23. Stern, Y. What is cognitive reserve? theory and research application of the reserve concept. J. international neuropsychological society 8, 448–460 (2002).

24. Raffegeau, T. E. et al. A meta-analysis: Parkinson’s disease and dual-task walking. Park. & Relat. Disord. 62, 28–35 (2019).

25. Yogev-Seligmann, G., Hausdorff, J. M. & Giladi, N. The role of executive function and attention in gait. Mov. disorders: official journal Mov. Disord. Soc. 23, 329–342 (2008).

26. Podsiadlo, D. & Richardson, S. The timed “up & go”: a test of basic functional mobility for frail elderly persons. J. Am. geriatrics Soc. 39, 142–148 (1991).

27. Mirelman, A. et al. Fall risk and gait in parkinson’s disease: the role of the lrrk2 g2019s mutation. Mov. Disord. 28, 1683–1690 (2013).

28. Herman, T., Weiss, A., Brozgol, M., Giladi, N. & Hausdorff, J. M. Gait and balance in parkinson’s disease subtypes: objective measures and classification considerations. J. neurology 261, 2401–2410 (2014).

29. Armstrong, M. J., Duff-Canning, S., Psych, C., Kowgier, M. & Marras, C. Independent application of montreal cognitive assessment/mini-mental state examination conversion. Mov. Disord. 30, 1710–1711 (2015).

30. Corrigan, J. D. & Hinkeldey, N. S. Relationships between parts a and b of the trail making test. J. clinical psychology 43, 402–409 (1987).

31. Martinez-Martin, P. et al. Expanded and independent validation of the movement disorder society–unified parkinson’s disease rating scale (mds-updrs). J. neurology 260, 228–236 (2013).

32. Robin, X. et al. proc: an open-source package for r and s+ to analyze and compare roc curves. BMC bioinformatics 12, 1–8 (2011).

33. Wilson, D. J. The harmonic mean p-value for combining dependent tests. Proc. Natl. Acad. Sci. 116, 1195–1200 (2019).

34. Lundberg, S. M., Erion, G. G. & Lee, S.-I. Consistent individualized feature attribution for tree ensembles. arXiv preprint arXiv:1802.03888 (2018).

