## Appendix A for "A novel performance scoring quantification framework for stress test set-ups"

### 1 Appendix: Derivatives

The suggested modified objective function derivative according to $x^d_{l,r}$ is:

$$\frac{\partial C}{\partial x^d_{l,r}} = \frac{1}{|\mathcal{B}|} \cdot \Bigg[ \sum_{\langle l,j \rangle \in \mathcal{B}} \frac{1}{|D_{lj}|} \cdot \frac{\partial L_h}{\partial \left(1 - \left(\bar{F}\left(x^d_{l,\cdot}\right) - \bar{F}\left(x^d_{j,\cdot}\right)\right)\right)} \cdot \left(-\frac{1}{\left|\mathcal{R}^d_l\right|}\right) +$$
$$\sum_{\langle i,l \rangle \in \mathcal{B}} \frac{1}{|D_{il}|} \cdot \frac{\partial L_h}{\partial \left(1 - \left(\bar{F}\left(x^d_{l,\cdot}\right) - \bar{F}\left(x^d_{j,\cdot}\right)\right)\right)} \cdot \left(\frac{1}{\left|\mathcal{R}^d_{l,\cdot}\right|}\right) \Bigg] +$$
$$\frac{\lambda_{\mathcal{W}}}{I} \cdot \frac{1}{|\mathcal{W}_l|} \cdot \Bigg[ \sum_{\langle d,d' \rangle \in D_i} \frac{\partial L_h}{\partial \left(1 - \left(F\left(x^d_{l,r}\right) - F\left(x^{d'}_{l,r}\right)\right)\right)} \cdot \left(-\frac{1}{\left|\mathcal{R}^d_l\right|}\right) +$$
$$\sum_{\langle d',d \rangle \in D_i} \frac{\partial L_h}{\partial \left(1 - \left(F\left(x^{d'}_{l,r}\right) - F\left(x^d_{l,r}\right)\right)\right)} \cdot \left(\frac{1}{\left|\mathcal{R}^d_l\right|}\right) \Bigg] +$$
$$\frac{\lambda_{\mathcal{R}}}{I} \cdot \frac{1}{|D_l|} \cdot \frac{2}{\left|\mathcal{R}^d_i\right|} \left[ \left(F\left(x^d_{l,r}\right) - \bar{F}\left(x^d_{l,\cdot}\right)\right)\left(1 - \frac{1}{\left|\mathcal{R}^d_l\right|}\right) + \sum_{\substack{r'=1 \\ r \neq r'}}^{R} \left(F\left(x^d_{l,r'}\right) - \bar{F}\left(x^d_{l,\cdot}\right)\right)\left(-\frac{1}{\left|x^d_{l,\cdot}\right|}\right)\right] \quad (1)$$

If we remove all terms equal to 0, then we are left with:

$$\frac{\partial C}{\partial x^d_{l,r}} = \frac{1}{|\mathcal{B}|} \cdot \Bigg[ \sum_{\langle l,j \rangle \in \mathcal{B}} \frac{1}{|D_{lj}|} \cdot \frac{\partial L_h}{\partial \left(1 - \left(\bar{F}\left(x^d_{l,\cdot}\right) - \bar{F}\left(x^d_{j,\cdot}\right)\right)\right)} \cdot \left(-\frac{1}{\left|\mathcal{R}^d_l\right|}\right) +$$
$$\sum_{\langle i,l \rangle \in \mathcal{B}} \frac{1}{|D_{il}|} \cdot \frac{\partial L_h}{\partial \left(1 - \left(\bar{F}\left(x^d_{l,\cdot}\right) - \bar{F}\left(x^d_{j,\cdot}\right)\right)\right)} \cdot \left(\frac{1}{\left|\mathcal{R}^d_l\right|}\right) \Bigg] +$$
$$\frac{\lambda_{\mathcal{W}}}{I} \cdot \frac{1}{|\mathcal{W}_l|} \cdot \Bigg[ \sum_{\langle d,d' \rangle \in D_i} \frac{\partial L_h}{\partial \left(1 - \left(F\left(x^d_{l,r}\right) - F\left(x^{d'}_{l,r}\right)\right)\right)} \cdot \left(-\frac{1}{\left|\mathcal{R}^d_l\right|}\right) +$$
$$\sum_{\langle d',d \rangle \in D_i} \frac{\partial L_h}{\partial \left(1 - \left(F\left(x^{d'}_{l,r}\right) - F\left(x^d_{l,r}\right)\right)\right)} \cdot \left(\frac{1}{\left|\mathcal{R}^d_l\right|}\right) \Bigg] +$$
$$\frac{\lambda_{\mathcal{R}}}{I} \cdot \frac{2}{|D_l|} \cdot \frac{\left(F\left(x^d_{l,r}\right) - \bar{F}\left(x^d_{l,\cdot}\right)\right)}{\left|\mathcal{R}^d_l\right|} \quad (2)$$

where $\frac{\partial L_h}{\partial c} = 0 \cdot \mathbb{1}_{(c<-h)} + \frac{(c+h)}{2h} \cdot \mathbb{1}_{(|c|<h)} + 1 \cdot \mathbb{1}_{(c>h)}$
